# Testing for the fitness benefits of natural transformation during community-embedded evolution

**DOI:** 10.1101/2023.03.20.532548

**Authors:** Macaulay Winter, Klaus Harms, Pål Jarle Johnsen, Angus Buckling, Michiel Vos

## Abstract

Natural transformation is a process where bacteria actively take up DNA from the environment and recombine it into their genome or reconvert it into extra-chromosomal genetic elements. The evolutionary benefits of transformation are still under debate. One main explanation is that foreign allele and gene uptake facilitates natural selection by increasing genetic variation, analogous to meiotic sex. However, previous experimental evolution studies comparing fitness gains of evolved transforming- and isogenic non-transforming strains have yielded mixed support for the “sex hypothesis.” Previous studies testing the sex hypothesis for natural transformation have largely ignored species interactions, which theory predicts provide conditions favourable to sex. To test for the adaptive benefits of bacterial transformation, the naturally transformable wildtype *Acinetobacter baylyi* and a transformation-deficient Δ*comA* mutant were evolved for five weeks. To provide strong and potentially fluctuating selection, *A. baylyi* was embedded in a community of five other bacterial species. DNA from a pool of different *Acinetobacter* strains was provided as a substrate for transformation. No effect of transformation ability on the fitness of evolved populations was found, with fitness increasing non-significantly in most treatments. Populations showed fitness improvement in their respective environments, with no apparent costs of adaptation to competing species. Despite the absence of fitness effects of transformation, wildtype populations evolved variable transformation frequencies that were slightly greater than their ancestor which potentially could be caused by genetic drift.

## Introduction

Natural transformation is a process whereby bacteria actively take up free DNA from the environment during a physiological state termed competence, followed by the recombination of this DNA into the recipient’s genome (or its reconversion into extra-chromosomal genetic elements). Natural transformation has been demonstrated in 80+ species across divergent bacterial lineages (Johnston *et al*., 2014) but is likely to be present in more species. Natural transformation can mediate the cell-to-cell transfer of large tracts of DNA, including virulence (Frosch and Meyer, 1992), antibiotic resistance (Blokesch, 2016; Winter *et al*., 2021) and metabolic genes (Tumen-Velasquez *et al*., 2018), making it one of the main prokaryote Horizontal Gene Transfer (HGT) mechanisms. The uptake of free DNA from the environment has been argued to provide three distinct (but not mutually exclusive) potential benefits to cells. First, as a source of nucleotides to be used for energy or building blocks (with recombination or maintenance of extrachromosomal DNA being a by-product) (Redfield, 1993, 2001; Hülter *et al*., 2017), second, to serve as templates for repairing genetic damage (Michod, Wojciechowski and Hoezler, 1988; Hoelzer and Michod, 1991; Mongold, 1992; Steinmoen, Knutsen and Håvarstein, 2002; Guiral *et al*., 2005; Ambur *et al*., 2016; Hülter *et al*., 2017), and third as a mechanism to create genetic variation (Vos, 2009; Ambur *et al*., 2016).

The genetic variation or ‘sex’ function of natural transformation has received most attention. Unlike many other HGT mechanisms, natural transformation is not mediated by Mobile Genetic Elements but is solely under the control of the recipient cell (Seitz and Blokesch, 2013; Johnston *et al*., 2014; Dubnau and Blokesch, 2019) and therefore could be assumed to be adaptive. By recombining adaptive alleles and genes in the same genomic background, natural transformation can result in the avoidance of clonal interference, allowing populations to adapt more rapidly. Indeed, both mathematical modelling (Levin and Cornejo, 2009; Engelstädter and Moradigaravand, 2013; Peabody, Li and Kao, 2017) and laboratory evolution experiments (Baltrus, Guillemin and Phillips, 2007; Woods *et al*., 2020; Nguyen *et al*., 2022) have supported this hypothesis. However, there remains controversy about the sex function of transformation, and not all experimental studies have found that fitness of recombining wildtype cells increased after evolution compared to isogenic, non-recombinogenic mutants. For instance, evolution experiments utilising the model system *Acinetobacter baylyi* found transformation-mediated fitness benefits either to be present (Perron *et al*., 2012), equivocal (Renda *et al*., 2015; Utnes *et al*., 2015) or absent (Bacher, Metzgar and De Crécy-Lagard, 2006; McLeman *et al*., 2016). Multiple studies found the ability to transform was lost during experimental evolution, indicating that any potential recombination-mediated fitness benefits were outweighed by the cost of maintaining the molecular machinery involved (Bacher, Metzgar and De Crécy-Lagard, 2006; Renda *et al*., 2015).

Studies to date lack interactions with multiple competitors that almost certainly characterise most natural situations. This could be an important shortcoming, as for sex to remain selectively advantageous it is necessary for selection to be strong and dynamic (Charlesworth, 1993; Burt, 2000), and interspecific competitors could greatly influence both these requirements (Otto and Nuismer, 2004; Kawecki *et al*., 2012). However, while interspecific competition can create fluctuating conditions, it can also constrain evolution (Luján *et al*., 2022). Experimental evolution approaches have hitherto relied on evolving recombining clones in isolation. To study the evolutionary benefits of natural transformation in the context of species interactions, we here experimentally evolve a transformable *A. baylyi* wildtype and a non-transformable isogenic Δ*comA* mutant for five weeks in the presence of other species (*i*.*e*., under biotic conditions) or alone (*i*.*e*., under abiotic conditions). Specifically, we use a system of five bacterial species which have been previously shown to stably coexist (Castledine, Padfield and Buckling, 2020; Padfield *et al*., 2020; Newbury *et al*., 2022). Our experimental evolution approach allows us to test 1) whether the evolved wildtype strain will be fitter than its non-recombining counterpart, specifically after evolution under biotic conditions and 2) whether transformation rate of the evolved wildtype has changed in response to these treatments.

## Material and Methods

### *Acinetobacter baylyi* ADP1 constructs

Two variants of the wildtype *A. baylyi* ADP1 strain with chromosomally encoded GFP and RFP, respectively were constructed using fluorescence::AMR cassettes derived from strains gifted by the Charpentier lab (Claude Bernard University, Lyon), and Hasty lab (University of San Diego, California), respectively. Briefly, the *sfGFP::apraR* cassette (Charpentier and Laaberki, unpublished) and *mCherry::specR* (Cooper, Tsimring and Hasty, 2017) cassettes were amplified via PCR with 1kb flanking regions homologous to the *att*Tn*7* locus and a putative prophage p4 region, respectively. Primers used for the *sfGFP::apraR* and *mCherry::specR* cassettes were 5’3 AAAGCCAATCGCTGACAGATGGTGG-3’, 5’-TTGGTCAGTGCCTGTCTTGCTGGTGAGCCGGTACGC-3’, and 5’-TCACCTGCATCCACTCAAGTGTCGTTT-3’, 5’-AAAGCCAATCGCTGACAGATGGTGG-3’, respectively (Integrated DNA Technologies, USA). PCR products were added to *A. baylyi* in LB Miller broth (Formedium, England) at 37°C and 180rpm for 24 hours at 1μg/mL to allow for chromosomal recombination of PCR amplificates via natural transformation Transformants were isolated by plating on LB agar containing 240μg/mL apramycin (Duchefa, The Netherlands), or 360μg/mL spectinomycin (Melford, UK), respectively. Next, non-competent counterparts of each fluorescent strain were generated by deletion of the *comA* gene via sequential natural transformation with linearised plasmids pKHNH6 and pKHNH3. Plasmids pKHNH6 and pKHNH3 were linearised prior to transformation using restriction enzymes EcoRV (Promega, USA) and KpnI (Fisher, USA), respectively. pKHNH6 carried a *comA*^*+*^ ::(*nptII sacB*) allele embedded in its natural flanking regions, and natural transformation of the respective fluorescence-marked *A. baylyi* strains by linearized plasmid DNA resulted in transformation-proficient isolates that were kanamycin-resistant and sensitive to 50g/L sucrose. Transformation of those respective isolates by pKHNH3 (carrying a Δ*comA* allele) resulted in a sucrose-resistant, kanamycin-susceptible and transformation-deficient strains, respectively Deletion of *comA* was verified using agar containing 50g/L sucrose, agar containing 10ug/mL kanamycin, and by PCR using primers 5’-TTGGTGTGATTGGTACGGTGGCTGGTGC-3’ and 5’-CTTGCAGACGATTGCTTACCTCAGCACTCGG-3’. The non-competent *A. baylyi* strains were confirmed to be non-transformable at the detectable limit (10-7) in all (6) technical replicates.

### Five-species community

The five-species community was composed of compost isolates belonging to the genera *Pseudomonas, Achromobacter, Variovorax, Ochrobactrum* and *Stenotrophomonas* (as identified by 16S rRNA sequencing) (Castledine, Padfield and Buckling, 2020; Padfield *et al*., 2020) (Table 1).

**Table 1.**
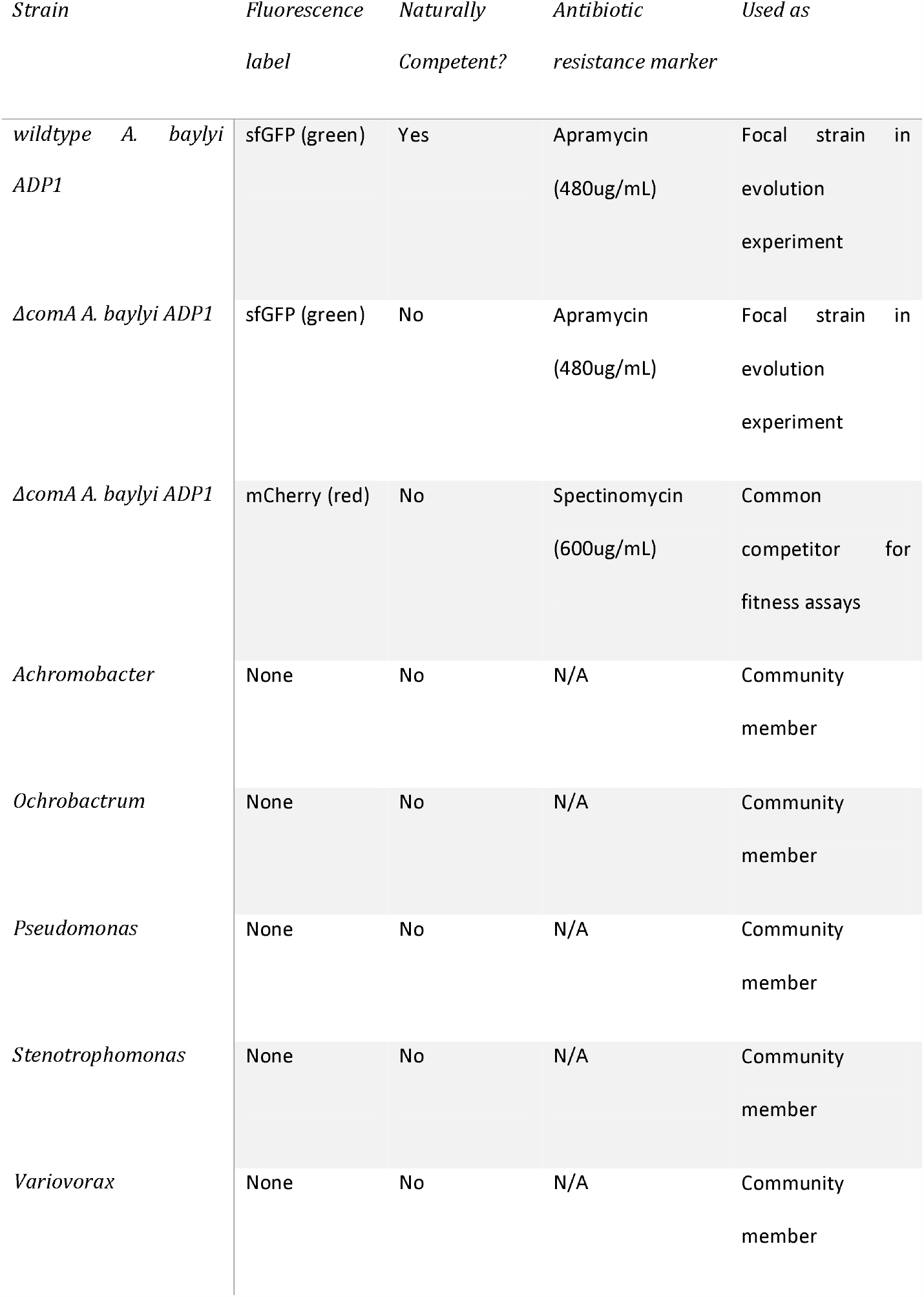
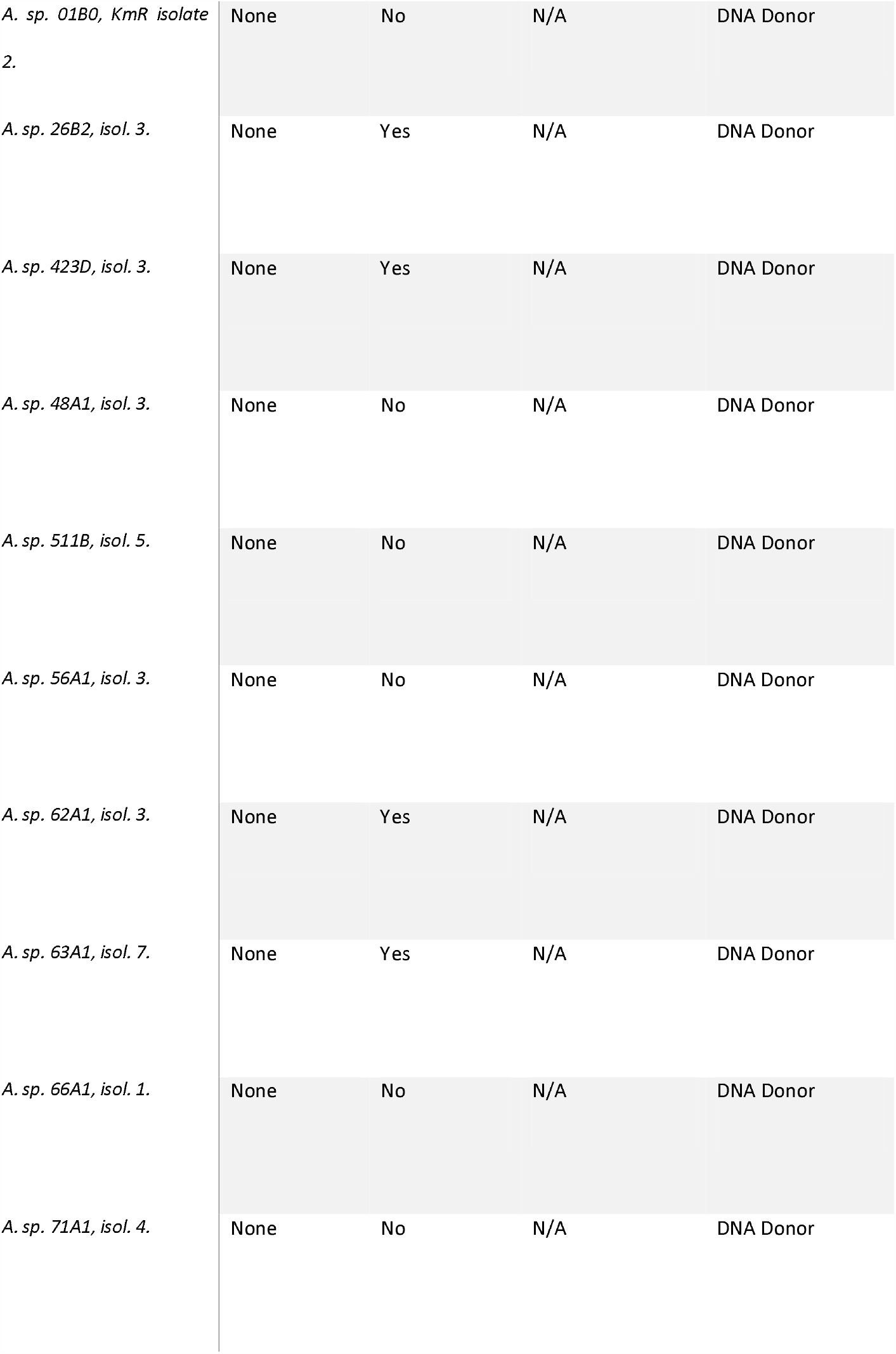
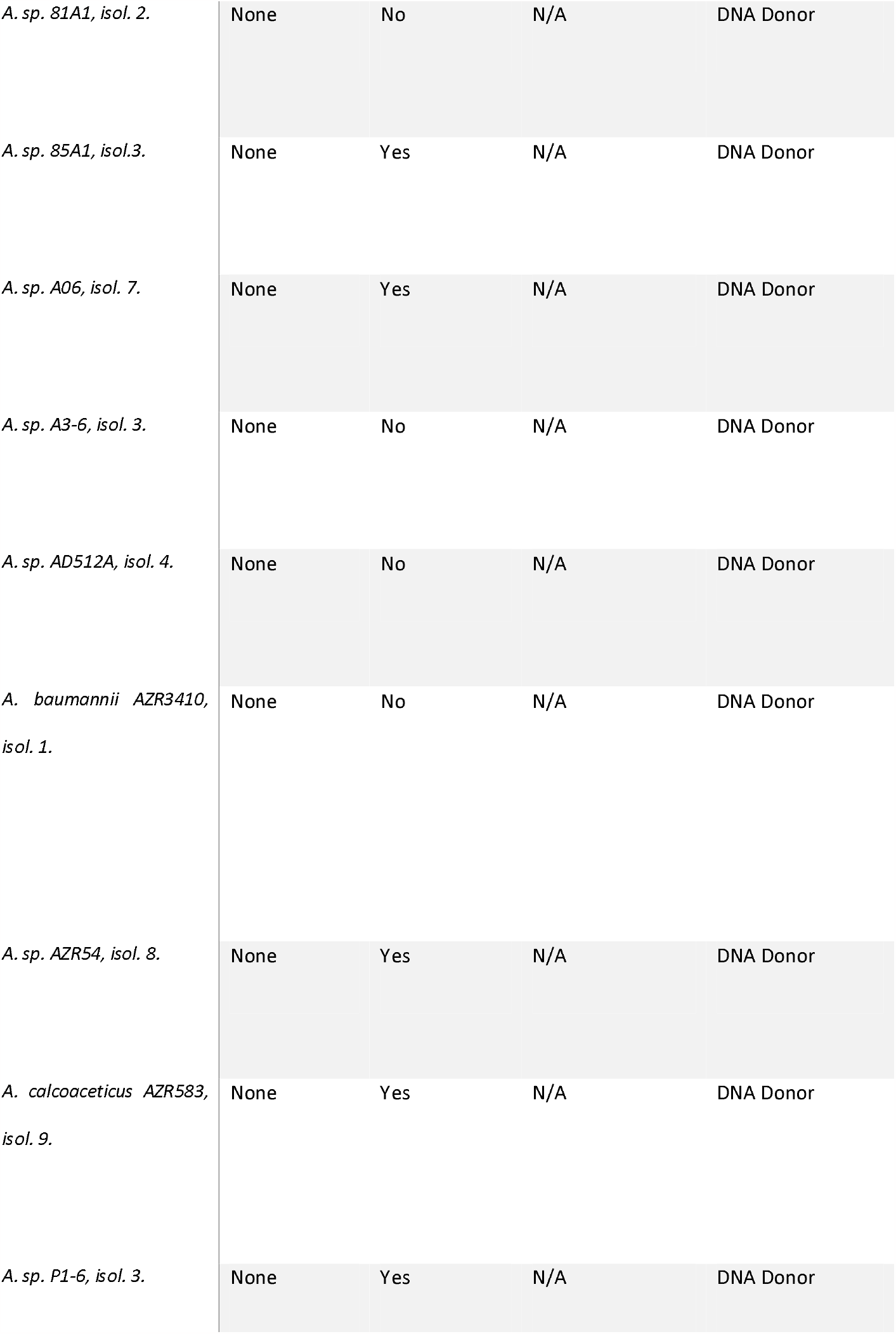
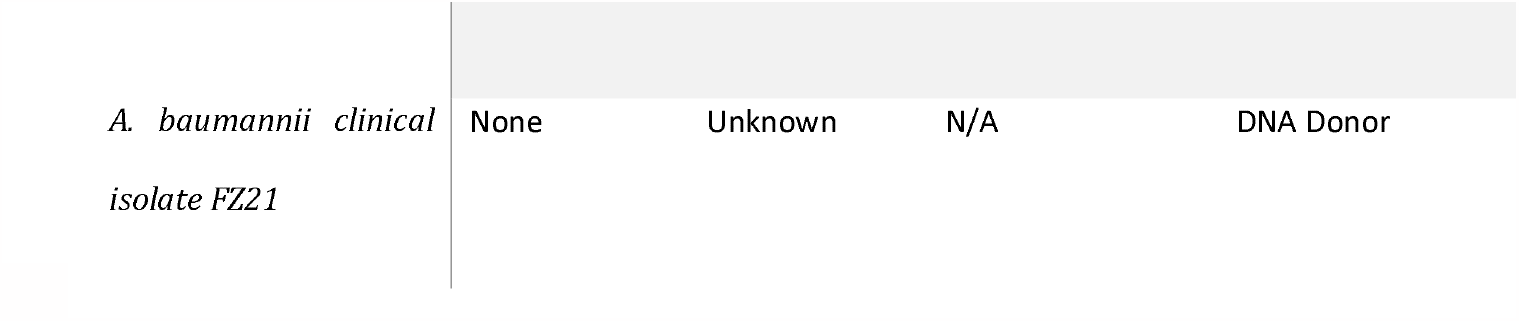
List of strains used in this study.

### Evolution experiment

The apramycin resistant wildtype and Δ*comA A. baylyi* ADP1 focal strains were separately propagated in three distinct competition environments with sixfold replication. Experimental treatments included competition against none, or all five of the five-species community concurrently (Figure 1). Cultures were grown in 25% TSB medium at 28°C in static conditions in glass microcosms with one layer of ColiRoller glass beads (Millipore, Merck, USA) covering the base of the microcosm to provide additional spatial structure. Instead of transferring a small volume to new microcosms, spent nutrient broth was replaced with fresh broth in the same microcosms, resulting in approximately 34-fold daily dilutions. This approach maintained spatial structure and saved on glass and plastic waste (Alves *et al*., 2021). Every 7^th^ day, cultures were vortexed, diluted 10-fold and plated on LB agar plates containing 240μg/ml apramycin to select for the green-fluorescent wildtype or Δ*comA A. baylyi* focal strain. After 24 hours of incubation at 37°C, the bacterial lawn from each individual replicate was scraped off with a sterile loop and transferred to a new microcosm for overnight growth (reaching stationary phase). Overnight cultures of the five competitor species were also made in LB broth at this time from –70°C freezer stocks. Equal volumes of overnight cultures of the focal *A. baylyi* strain and competitor species (where appropriate; Table 1) totalling 100uL was added to a new microcosm containing 9.9mL 25% TSB, commencing the next week of transfers. DNA as a substrate for transformation sourced from a pool of 20 *Acinetobacter* strains was added at the point of nutrient replenishment each day for each treatment (see next section). At the end of the final week of transfers, cultures were plated on LB agar supplemented with 240μg/ml apramycin where 100 colonies were picked and pooled for overnight growth and later frozen at -70°C in 25% glycerol prior to use in subsequent competition and transformation assays (Fig. 1).

**Figure 1.**
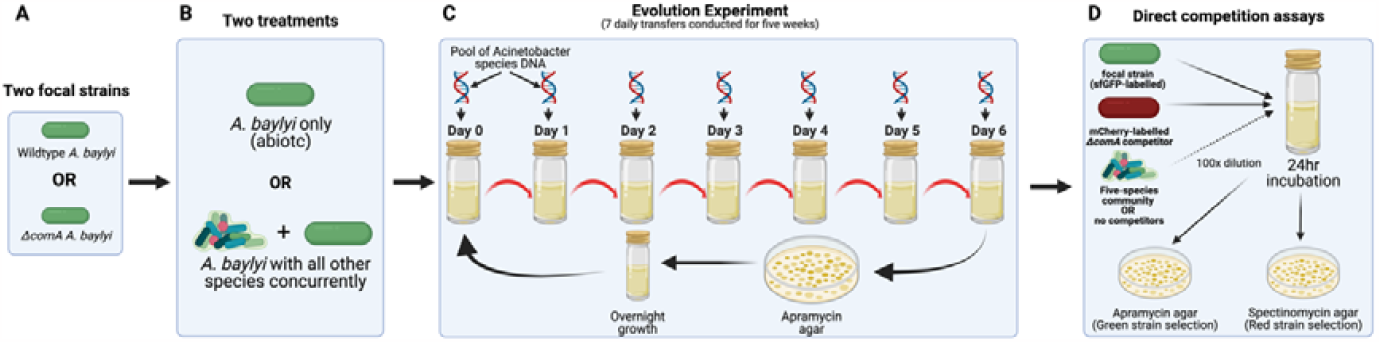
Illustration of the evolution experiment and direct competition assays. A: Two focal strains (competent wildtype and isogenic non-competent counterpart) were cultured separately in their respective treatment conditions (B). B: Focal strains were subjected to two treatment conditions: monoculture (abiotic) or co-culture with five competitor species. C: Cultures were propagated by replacing spent broth with fresh media resulting in an approximately 34-fold dilution each passage. After the 6^th^ passage, all species except the focal strain are killed off using LB agar amended with apramycin. The focal strain and freezer stocks of the competitors were allowed to grow to maximum density in LB broth before being inoculated together for another week of passaging. D: After five full weeks of passaging, the focal strains are selected for again with use of apramycin agar and frozen in 25% glycerol at –70°C until tested against a common competitor to measure relative fitness. Figure created with BioRender.com.

### Donor DNA

Twenty *Acinetobacter* strains (table 1; Ray and Nielsen, 2005; Alseth et al., 2019) were cultured separately in LB broth overnight. Cultures were then pooled in equal volumes and lysed following the Qiagen® Genomic DNA Handbook (April 2012) protocol. Combined DNA from the eluate was precipitated by adding two volumes of ice-cold ethanol and centrifuged at 26000xg for 15 minutes to pellet the DNA. DNA was dissolved in TE buffer to a final concentration of 227.2ng/μL (Nanodrop 2000, Thermo Scientific) by heating at 50°C and stirring for 16 hours. DNA was frozen at -20°C in single-use aliquots for addition to each daily transfer. When used during the evolution period, DNA was diluted to a final concentration of 250ng/ml (the saturating concentration of genomic DNA for transformation in *A. baylyi* (Overballe-Petersen *et al*., 2013)). The raw and annotated DNA sequencing data for the 19 strains were deposited at the European Nucleotide Archive under BioProject accession number PRJEB55833 (Winter *et al*., 2023).

### Competition assays

For all replicates of the five concurrent species community treatment and the abiotic treatment, 100 evolved green-fluorescent wildtype and Δ*comA A. baylyi* clones were picked, pooled and frozen at – 80°C before use. A mixture containing equal volumes of the green-fluorescent *A. baylyi* pool, a Δ*comA* red-fluorescent *A. baylyi*, and members of the five species community (where appropriate) was produced for each replicate. One hundred microlitres of mixed culture was immediately inoculated to 9.9ml of 25% TSB and grown for 24 hours in glass microcosms at 28°C with no agitation. Each of the six replicate evolved populations per treatment were competed with the differentially marked Δ*comA* strain in conditions identical to treatments in the evolution experiment (i.e., containing either all or none of the five competitor species). Samples were plated 0 hours and 24 hours after inoculation on LB agar supplemented with either apramycin (240μg/ml) or spectinomycin (360μg/mL) to determine the densities of the evolved (green)- and Δ*comA* (red)-fluorescent *A. baylyi*, respectively (Fig. 1).

### Transformation Assays

Red fluorescent, spectinomycin-resistance conferring marker DNA was obtained by heating overnight cultures of the *A. baylyi* ADP1 *mCherry::specR* at 70°C for 1.5 hours and centrifuging at 2500xg for 15 minutes and resuspending in reduced volume to produce a 50x concentration of lysate. Lysate was stored at -20°C for up to one month before downstream application. Freezer stocks of 100 pooled clones for each endpoint population were inoculated in LB broth and grown overnight at 37°C. Cultures were diluted ten-fold with LB broth supplemented with cell lysate at a final concentration of 2.5× maximum cell density and incubated for 3 hours at 37°C and 180 rpm. Cells were then plated on LB agar supplemented with 240μg/ml apramycin and 360μg/ml spectinomycin, and non-selective LB agar and incubated for 48 hours at 28°C. Transformation frequency was calculated by dividing the CFU/mL of the transformed (doubly fluorescent and dually-AMR) population by the total population CFU/mL. As a control for spontaneous spectinomycin resistance mutations, we included a treatment where no *mCherry::specR* DNA was added.

### Statistical Analysis

Normal distribution of the data was verified using Shapiro-Wilk tests. To determine the relative fitness of evolved lines relative to the unevolved control in competition experiments, the selection-rate constant was calculated as described in (Lenski *et al*., 1991). Selection-rate constant estimates were analysed with paired t-tests for comparisons using the mean value for each measured population (treatments tested within assay conditions and grouped by genotype and evolution treatment conditions). Because relative fitness values were often negative (i.e., the common competitor displayed greater fitness than the evolved focal strains), analyses in this assay were conducted using the selection-rate constant in lieu of the relative fitness parameter (Lenski *et al*., 1991). As ancestral populations were not significantly different to each other in fitness as a function of genotype (paired t-tests; biotic assay conditions, p=0.35, abiotic assay conditions, p=0.74), fitness measurements of the evolved populations were standardised to ancestors by subtracting the ancestral selection-rate constant (averaged for both genotypes) from that of the evolved population in all analyses. Standardised selection rates significantly different to 0 demonstrate fitness change.

Generalised statements about treatment effects observed in fitness assays were made using linear mixed effect models with the lme4 package v1.29 (Bolker, 2022). Model selection was achieved using backward stepwise regression, using biological replicates as a random explanatory variable. Model residuals were checked using the DHARMa package v0.4.5 (Hartig, 2022). All biological replicates for the selection-rate constant analyses were measured in triplicate and averaged before downstream analyses. Transformation frequencies were analysed non-parametrically with Wilcoxon tests as the data were not normally distributed (Shapiro-Wilks tests, p<0.001). T-tests, Wilcoxon, Kruskal-Wallis and Levene testing was conducted using the rstatix package v0.7.0 (Kassambara, 2021). In all analyses, a *p* value of less than 0.05 was considered significant. Multiple testing was corrected for with false discovery rate (FDR) correction.

## Results

### Natural transformation does not provide fitness benefits in a community context

To test whether natural transformation favours adaptation in a biotic community context compared to an abiotic environment, we evolved a recombining wildtype *A. baylyi* and an isogenic non-recombining Δ*comA* mutant supplemented with DNA extracted from a pool of conspecifics as a substrate for natural transformation. This five species community has been shown to stably coexist in 1/64^th^ strength tryptic soy broth (TSB), and 28°C with weekly passages (Padfield *et al*., 2020), but was found to also exist with *A. baylyi* stably in 25% TSB medium with daily transfers (results not shown). After five weeks of evolution, changes in fitness of each evolved line relative to a single unevolved, differentially marked Δ*comA* strain were measured using pairwise competition assays. Populations were assayed in both presence and absence of competitors. All populations increased in fitness relative to ancestral populations (1 sample t-tests: p<0.01, corrected for multiple testing, Figure 2, Table 2), except for the wildtype strains evolved in abiotic and biotic conditions when assayed in biotic conditions (p=0.486, and p=0.058, respectively; Figure 2, Table 2).

**Figure 2.**
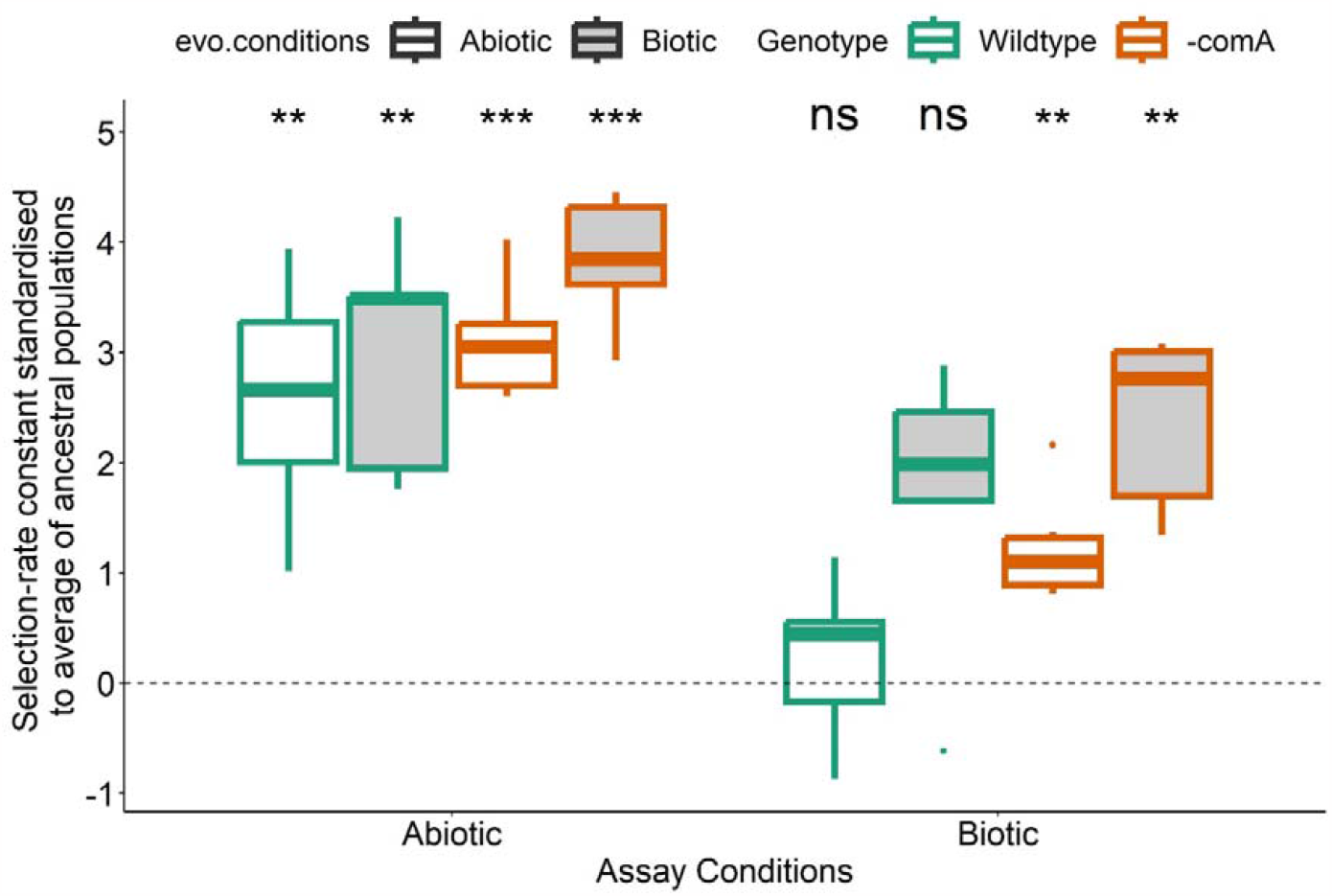
Selection-rate constants of evolved populations standardised to ancestral populations in respective assay conditions (y=0). Bracketed asterisks describe significant differences (t-test) between evolved populations within the same assay conditions. Non-bracketed asterisks describe significant differences (t-test) between evolved populations’ and 0. Fitness differences of each of the six biological replicates per treatment were measured in triplicate. (ns, no significant difference; *, p<0.05; **, p<0.01; ***, p<0.001).

Linear mixed effects models revealed no interactions between explanatory variables: genotype, evolutionary conditions, and assay conditions when predicting fitness increases (X^2^ = 4.2241, df =4, p = 0.3765; Figure 2). Genotype and evolution conditions are very significant explanatory variables for model predictions (p<0.001 and p<0.01, respectively). The Δ*comA* populations appear better adapted to the experimental environments relative to the wildtype (Welch’s two sample t-test, p=0.4401; Figure 2). Adaptation to abiotic conditions occurs more effectively than to biotic conditions, irrespective of evolution conditions or genotype (t-test, p<0.0001). Evolution in a biotic environment leads to greater adaptation to biotic environments, but no greater adaptation to abiotic environments (pairwise t-testing, p<0.01, and p=0.113, respectively; Figure 2). Pairwise t-tests of the differences in selection-rate constants of the evolved populations (grouped by genotype and evolution conditions) tested separately with respect to their assay conditions (biotic and abiotic) showed no significant differences (p>0.05; Figure 2; Table 3).

**Table 2.**
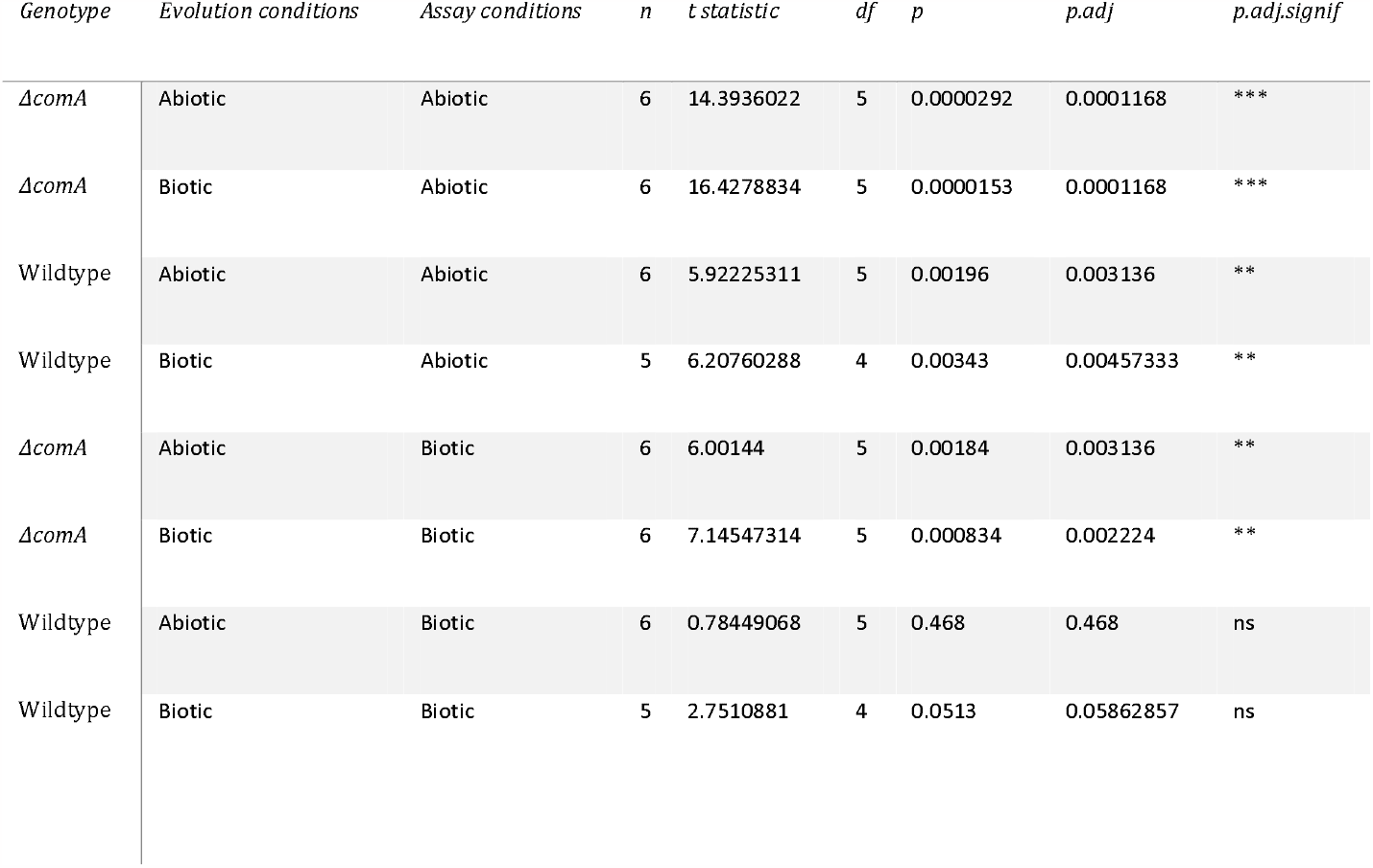
Pairwise comparisons of evolved populations’ fitness gains after 5 weeks of evolution compared to a differentially marked Δ*comA* A. baylyi strain (t-test).

| *Genotype* | *Evolution conditions* | *Assay conditions* | *n* | *t statistic* | *df* | *p* | *p.adj* | *p.adj.signif* |
| --- | --- | --- | --- | --- | --- | --- | --- | --- |
| $\Delta comA$ | Abiotic | Abiotic | 6 | 14.3936022 | 5 | 0.0000292 | 0.0001168 | *** |
| $\Delta comA$ | Biotic | Abiotic | 6 | 16.4278834 | 5 | 0.0000153 | 0.0001168 | *** |
| Wildtype | Abiotic | Abiotic | 6 | 5.92225311 | 5 | 0.00196 | 0.003136 | ** |
| Wildtype | Biotic | Abiotic | 5 | 6.20760288 | 4 | 0.00343 | 0.00457333 | ** |
| $\Delta comA$ | Abiotic | Biotic | 6 | 6.00144 | 5 | 0.00184 | 0.003136 | ** |
| $\Delta comA$ | Biotic | Biotic | 6 | 7.14547314 | 5 | 0.000834 | 0.002224 | ** |
| Wildtype | Abiotic | Biotic | 6 | 0.78449068 | 5 | 0.468 | 0.468 | ns |
| Wildtype | Biotic | Biotic | 5 | 2.7510881 | 4 | 0.0513 | 0.05862857 | ns |

**Table 3.**
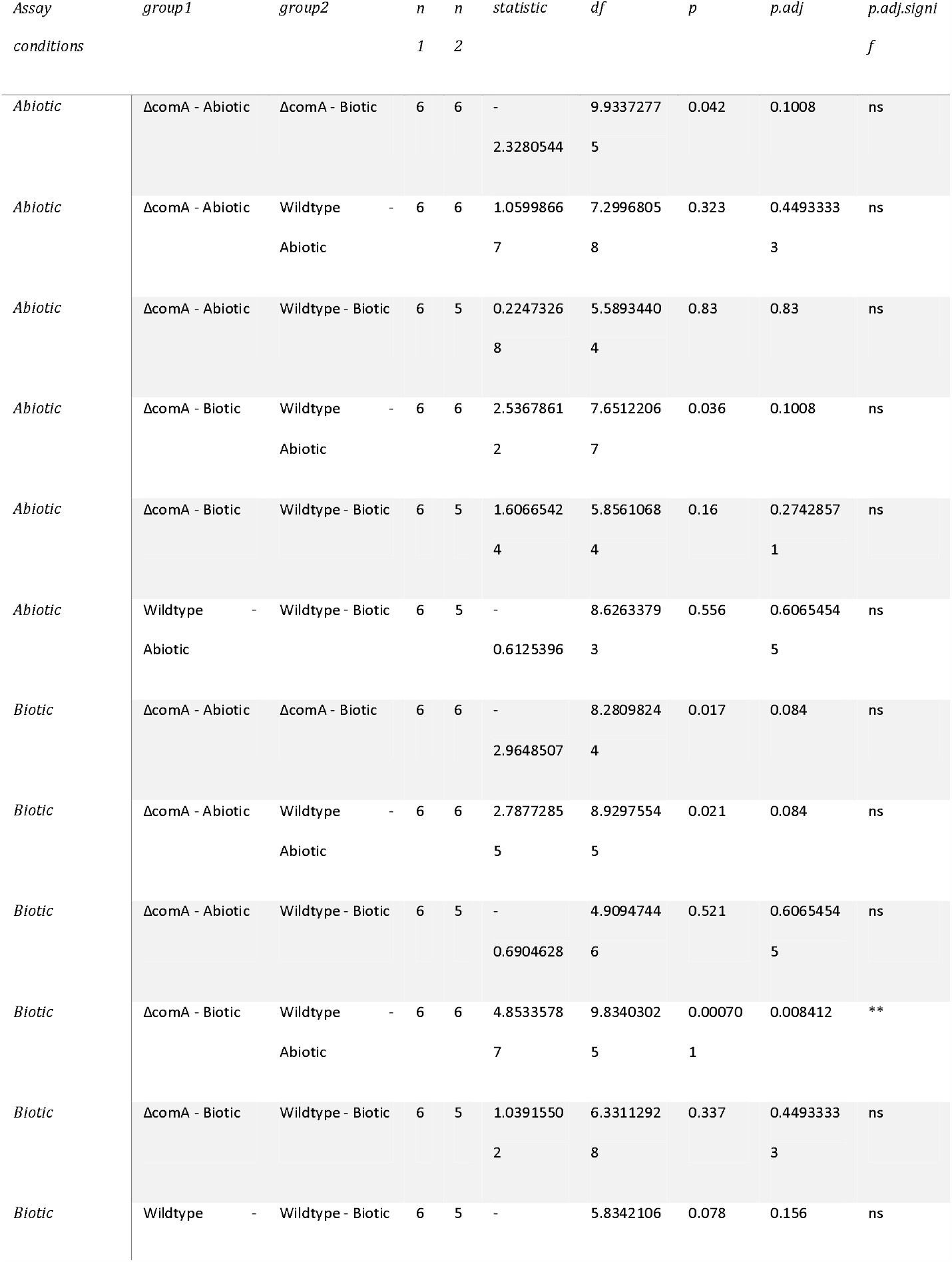
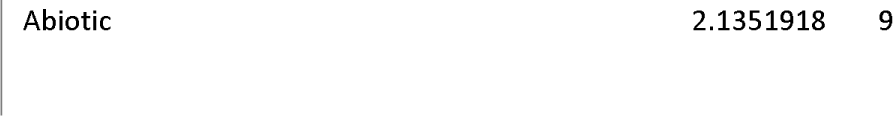
Pairwise comparisons of evolved populations’ fitness gains after 5 weeks of evolution (t-test).

### Transformation frequency did not decrease after evolution

To test whether the lack of fitness difference of the wildtype strain stemmed from a possible loss of natural transformation, we conducted transformation assays for the abiotic treatment and the five species community treatment (Figure 1). No significant differences in transformation frequency were found between the evolved wildtype lineages across the two evolution environments or the ancestor (Wilcoxon test, p>0.05; Figure 3; Table 4). While transformation frequency did not change significantly, observed transformation frequencies were higher in all evolved wildtype populations than in the ancestor. A separate experiment using donor DNA acquired through chemical lysis (the method used in the evolution experiment), resulted in a much lower transformation frequency for all tested samples and did not show a clear significant difference in transformation frequency between any test groups (p>0.05; Supplemental Figure 1; Table 5).

**Table 4.** Pairwise comparisons of evolved populations’ transformation frequency gains after 5 weeks of evolution (Wilcoxon-test). Samples are tested using heat-killed cell lysate.

| *group1* | *group2* | *n1* | *n2* | *statistic* | *p* | *p.adj* | *p.adj.signif* |
| --- | --- | --- | --- | --- | --- | --- | --- |
| *ΔcomA Ancestor* | Abiotic Culture | 1 | 6 | 0 | 0.286 | 0.4995 | ns |
| *ΔcomA Ancestor* | Community | 1 | 5 | 0 | 0.333 | 0.4995 | ns |
| *ΔcomA Ancestor* | WT Ancestor | 1 | 1 | 0 | 1 | 1 | ns |
| *Abiotic Culture* | Community | 6 | 5 | 16 | 0.931 | 1 | ns |
| *Abiotic Culture* | WT Ancestor | 6 | 1 | 6 | 0.286 | 0.4995 | ns |
| *Community* | WT Ancestor | 5 | 1 | 5 | 0.333 | 0.4995 | ns |

**Table 5.** Pairwise comparisons of evolved populations’ transformation frequency gains after 5 weeks of evolution (Wilcoxon-test). Samples are tested using chemically acquired cell lysate.

| *group1* | *group2* | *n1* | *n2* | *statistic* | *p* | *p.adj* | *p.adj.signif* |
| --- | --- | --- | --- | --- | --- | --- | --- |
| *ΔcomA Ancestor* | Abiotic Culture | 1 | 6 | 0.5 | 0.313 | 0.666 | ns |
| *ΔcomA Ancestor* | Community | 1 | 5 | 0 | 0.333 | 0.666 | ns |
| *ΔcomA Ancestor* | WT Ancestor | 1 | 1 | 0 | 1 | 1 | ns |
| *Abiotic Culture* | Community | 6 | 5 | 11 | 0.537 | 0.8055 | ns |
| *Abiotic Culture* | WT Ancestor | 6 | 1 | 4 | 0.857 | 1 | ns |
| *Community* | WT Ancestor | 5 | 1 | 5 | 0.333 | 0.666 | ns |

**Figure 3.**
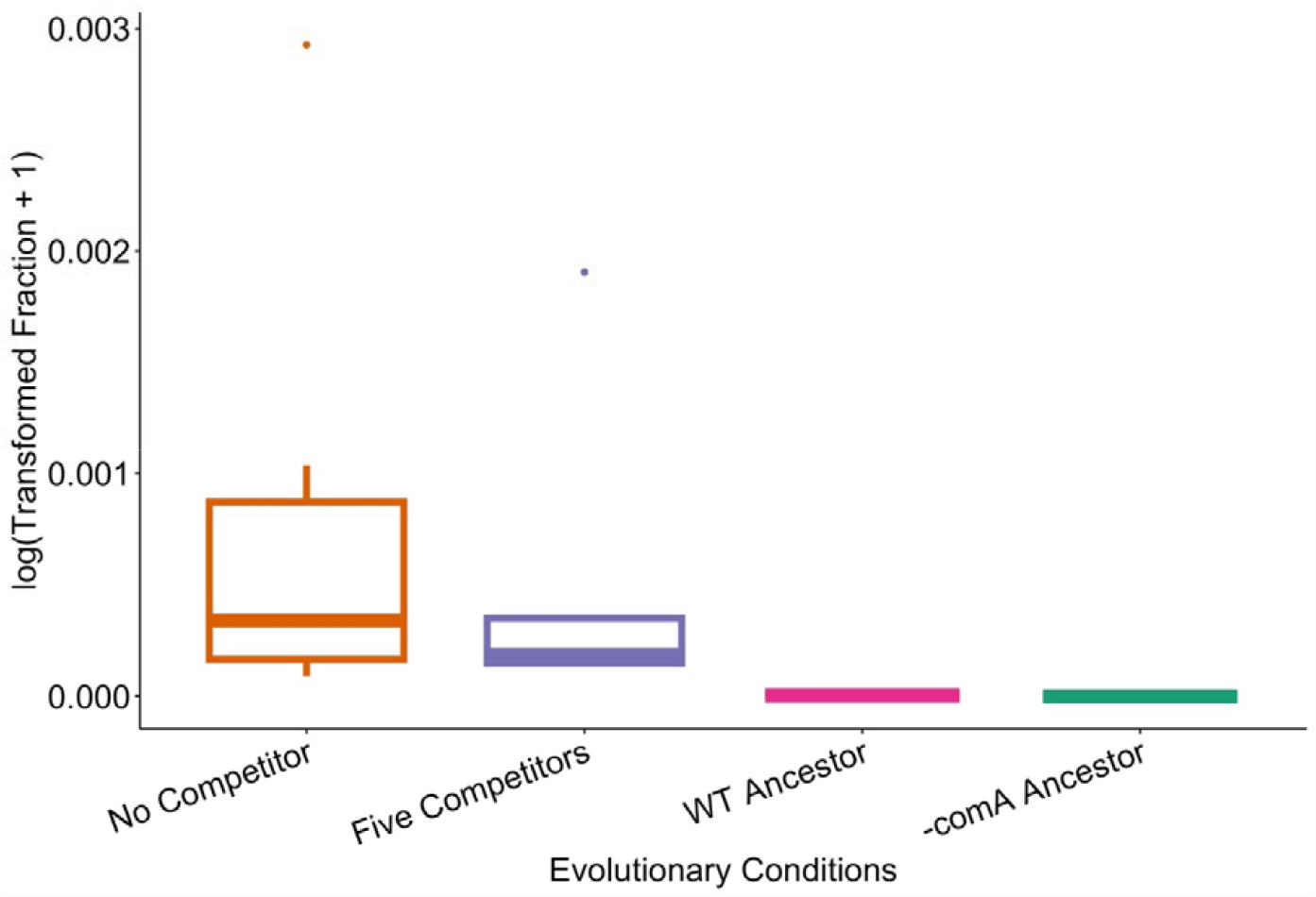
Treatment-level transformation frequencies of ancestral and evolved populations using heat-killed cell lysate. Transformation frequencies were measured in triplicate per biological replicate. The No Competitor, Five Competitor, WT ancestor, and Δ*comA* ancestor treatments had 6, 5, 1, and 1 biological replicates, respectively.

## Discussion

Here we tested if recombination mediated by natural transformation in *A. baylyi* provided adaptive benefits under biotic and abiotic experimental conditions. All replicate populations in either treatment increased fitness compared to their ancestors when assayed in abiotic conditions (Figure 2). Populations of the non-recombining (Δ*comA)* genotype also show increased fitness relative to their ancestor when assayed under biotic conditions regardless of evolutionary conditions. This contrasts with the recombining wildtype, which does not show significant increases in fitness after evolution in either environment, contrary to our hypothesis that sex is advantageous in the presence of other species. However, the absence of significant fitness increases observed in the wildtype which was evolved and tested in biotic conditions are probably due to a single tested population which is less fit than the ancestor, lowering the estimation of the mean (Figure 2). Further, no significant differences in fitness between evolved linages were found when comparing across genotypes and assaying in biotic conditions. Therefore, this study has not found a clear and significant beneficial (or deleterious) effect of natural transformation on adaptative evolution.

There is clear trade-off asymmetry displayed by evolving the wildtype *A. baylyi* in the presence or absence of the five competitor species (biotic and abiotic conditions, respectively). Evolution of the wildtype under biotic conditions did not constrain adaptation to an abiotic environment, but the converse is not true (Figure 2). This is because the factors in abiotic conditions (growth media, temperature, oxygen availability, spatial structure) were also consistent in the biotic condition and are therefore selected for in the biotic environment. However, there may be some adaptive mutations that arise under biotic conditions which are not advantageous in the abiotic environment, and therefore were not selected for during evolution in abiotic conditions (Gomez *et al*., 2022). Our data show that evolution in a biotic environment best prepares a population for a biotic environment, and does not constrain adaptation to an abiotic environment, regardless of the population’s ability to naturally transform.

Interestingly, mean transformation frequencies of all tested populations increased relative to the transformable ancestor, although this was not statistically significant. This contrasts with previous studies where transformation frequencies declined markedly after experimental evolution (Bacher, Metzgar and De Crécy-Lagard, 2006; Utnes *et al*., 2015). The maintenance, and potential increase of transformability in all evolved populations suggests that the ability to transform was not selected against and may have increased because of genetic drift, given that it served no apparent adaptive benefit. The increase of transformation frequency at varying rates per population (Figure 4) suggests that significant differences both within and between treatment groups may occur after a more prolonged evolution period.

**Figure 4.**
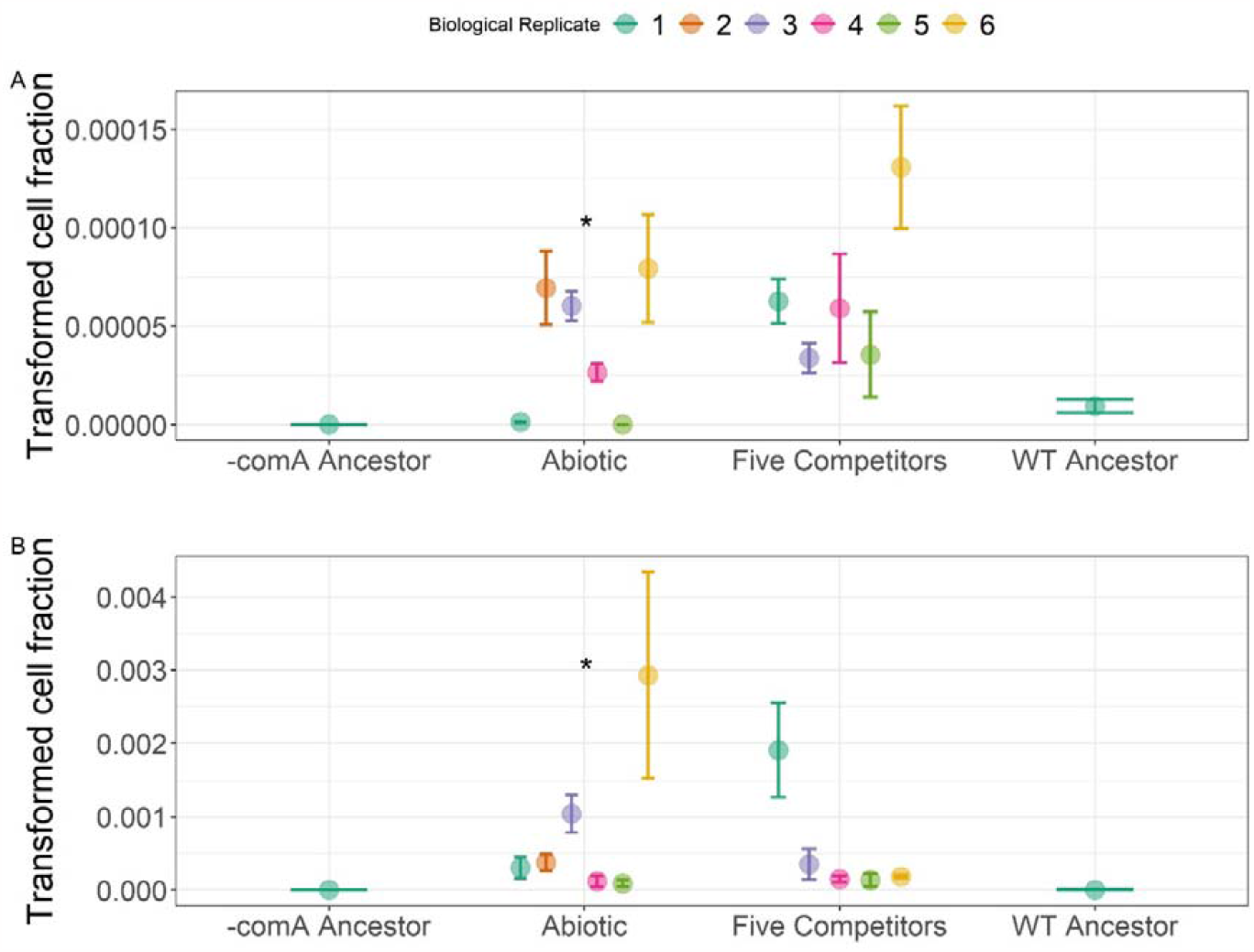
Variation of transformation frequencies of biological replicates within evolutionary treatment groups for (A) chemically acquired and (B) heat-killed cell lysate. Transformation frequencies were measured in triplicate per biological replicate. Points are biological replicates and bars are means ± one standard error. Asterisks denote significantly different transformation frequencies between intra-treatment biological replicates (Kruskal-Wallis, p<0.05).

Taken together, these findings show no significant evolutionary costs or benefits of transformation in *A. baylyi*. This does not mean that no bacterial species use transformation to facilitate faster adaptation, as the physiological contexts and molecular mechanisms of natural transformation vary across species (Johnston *et al*., 2014). Specifically, there is convincing evidence supporting the sex hypothesis for transformation in *H. pylori* (Baltrus, Guillemin and Phillips, 2007; Woods *et al*., 2020; Nguyen *et al*., 2022). There is no doubt that transformation can give an immediate adaptive benefit to *A. baylyi* such as in the contexts of antimicrobial resistance acquisition (Domingues *et al*., 2012; Perron *et al*., 2012; Hülter *et al*., 2017; Mantilla-Calderon *et al*., 2019), but its long-term benefits remain uncertain. It is possible that different growth conditions, interactions with live DNA donors, and reduced doubling rates of cells (limiting the frequency and contribution of mutation for adaptation), can better demonstrate the adaptive benefit of natural transformation in *A. baylyi* than what has been shown in this study.

## Acknowledgments

We thank the Charpentier lab (Claude Bernard University, Lyon) and Hasty lab (University of San Diego, California) for donation of *A. baylyi* strains. MW acknowledges support from the National Environment Research Council (NERC) and the GW4+ Doctoral Training Partnership (grant NE/S007504/1). MV acknowledges support from the National Environment Research Council (NERC; grant NE/T008083/1). KH and PJ acknowledge support from The Research Council of Norway (NFR; grant number 275672). We thank Daniel Padfield and Elze Hesse for help with statistical analyses.

The authors have no conflicts of interest to declare.

## Footnote

One treatment sample containing the wildtype strain in the presence of five concurrent competitors was lost near the end of the experimental evolution period. In the interest of ensuring all samples were directly comparable by experiencing identical conditions, we decided to not replace the lost sample by evolving it independently to the other samples. This explains why the replication level for one treatment is 5 instead of 6.

