## Supplemental Figure 1 for "Testing for the fitness benefits of natural transformation during community-embedded evolution"

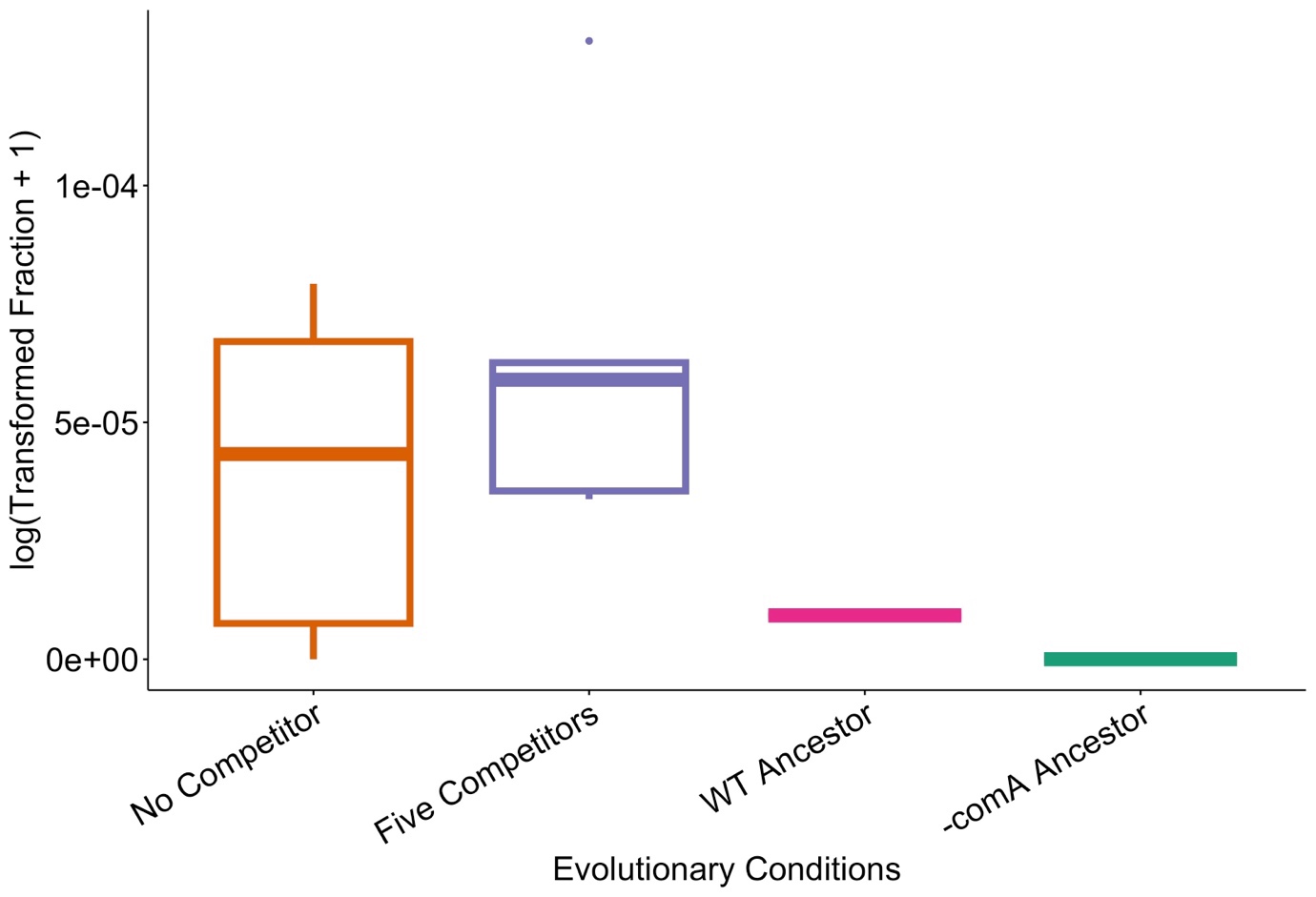

Supplemental Figure 1. Treatment-level transformation frequencies of ancestral and evolved populations using chemically acquired cell lysate. Transformation frequencies were measured in triplicate per biological replicate. The No Competitor, Five Competitor, WT ancestor, and ∆comA ancestor treatments had 6, 5, 1, and 1 biological replicates, respectively.
